# Meta-MultiSKAT: Multiple phenotype meta-analysis for region-based association test

**DOI:** 10.1101/593814

**Authors:** Diptavo Dutta, Sarah A. Gagliano Taliun, Joshua S. Weinstock, Matthew Zawistowski, Carlo Sidore, Lars G. Fritsche, Francesco Cucca, David Schlessinger, Gonçalo R. Abecasis, Chad M. Brummett, Seunggeun Lee

## Abstract

The power of genetic association analyses can be increased by jointly meta-analyzing multiple correlated phenotypes. Here, we develop a meta-analysis framework, Meta-MultiSKAT, that uses summary statistics to test for association between multiple continuous phenotypes and variants in a region of interest. Our approach models the heterogeneity of effects between studies through a kernel matrix and performs a variance component test for association. Using a genotype kernel, our approach can test for rare-variants and the combined effects of both common and rare-variants. To achieve robust power, within Meta-MultiSKAT, we developed fast and accurate omnibus tests combining different models of genetic effects, functional genomic annotations, multiple correlated phenotypes and heterogeneity across studies. Additionally, Meta-MultiSKAT accommodates situations where studies do not share exactly the same set of phenotypes or have differing correlation patterns among the phenotypes. Simulation studies confirm that Meta-MultiSKAT can maintain type-I error rate at exome-wide level of 2.5×10^−6^. Further simulations under different models of association show that Meta-MultiSKAT can improve power of detection from 23% to 38% on average over single phenotype-based meta-analysis approaches. We demonstrate the utility and improved power of Meta-MultiSKAT in the meta-analyses of four white blood cell subtype traits from the Michigan Genomics Initiative (MGI) and SardiNIA studies.

## Introduction

The advent of large scale genome-wide association studies (GWAS) has shown that many distinct phenotypes share substantial genetic etiology(Bulik-Sullivan et al., 2015) and many loci have pleiotropic effects(Cotsapas et al., 2011; Purcell, Smoller, Cotsapas, Solovieff, & Lee, 2013; Sivakumaran et al., 2011). To leverage the widespread pleiotropy, a statistical model to jointly test multiple phenotypes is beneficial. Although data on multiple related phenotypes are often collected in hospital or population based studies, association tests are usually performed with one phenotype at a time. Such methods that do not account for the correlation between phenotypes may lack power to detect cross-phenotype effects of associated loci(Ferreira & Purcell, 2009; Huang, Johnson, & O’Donnell, 2011; Ray, Pankow, & Basu, 2016). Alternatively, joint tests which aggregate association signals in multiple phenotypes can substantially improve power over single phenotype-based tests(Ferreira & Purcell, 2009; Ray et al., 2016; Ried et al., 2012; Zhou & Stephens, 2014).

Meta-analysis of multiple studies, using association summary statistics, is a practical approach to increase power by increasing sample sizes(Panagiotou, Willer, Hirschhorn, & Ioannidis, 2013). Meta-analysis is especially valuable for association analysis of variation on the lower end of the allele frequency spectrum, since detecting such associations often require large sample sizes. It seems logical to expect that meta-analyzing multiple phenotypes can further increase power of rare variant tests. Various methods have been developed for meta-analysis of multiple phenotypes(Majumdar, Haldar, Bhattacharya, & Witte, 2018; Ray & Boehnke, 2018; Zhu et al., 2015), but most of them are single variant-based methods, which have low power to identify rare variant associations. More powerful gene or region-based tests for multiple phenotypes have been developed for use within a single study(Broadaway et al., 2016; Selyeong Lee et al., 2017; B. Wu & Pankow, 2016). However, to the best of our knowledge, no work has been done to extend these methods to meta-analysis. This is partly because most of the methods are similarity-based non-parametric methods, which are difficult to extend to meta-analysis. We have developed a regression-based method, Multiple phenotype sequence kernel association test (Multi-SKAT)(Dutta, Scott, Boehnke, & Lee, 2019), that can aggregate signals across models with different kernels, which cannot be done by current methods.

In this article we propose Meta-MultiSKAT, a meta-analysis extension of Multi-SKAT, which uses summary statistics. Meta-MultiSKAT models the relationship between effect sizes of different studies through a kernel matrix and performs a variance component test of association. Our method is based on summary statistics from individual studies and retains useful features of Multi-SKAT, including fast computation. Meta-MultiSKAT can incorporate various missing data scenarios, including situations where studies do not share exactly the same set of phenotypes, and test for only rare variants as well as for the combined effects of both common and rare variants. The latter allows us to evaluate the overall effect of gene or region on multiple phenotypes. By using kinship adjusted score statistics, Meta-MultiSKAT can account for sample relatedness, an important feature to use in a study with widespread relatedness, such as the SardiNIA study(Sidore et al., 2015; Vacca et al., 2006). To avoid loss of the power due to model misspecification, we have also developed a minimum p-value-based omnibus test that can aggregate results across different patterns of association. We evaluate the performance of our method through extensive type-I error and power simulations.

We applied Meta-MultiSKAT to meta-analyze four white blood cell (WBC) subtype traits from the Michigan Genomics Initiative (MGI)(Fritsche et al., 2018) study and the SardiNIA study(Sidore et al., 2015; Vacca et al., 2006). In addition to detecting the genes *PRG2* [MIM: 605601] and *RP11-872D17.8*, that had significant association signals with WBCs within one of the studies, Meta-MultiSKAT further identified two additionally associated genes (*IRF8* [MIM: 601565] and *CCL24* [MIM: 602495]) that did not have any significant signals in either of the studies but were identified as significant only as a result of meta-analysis.

## Material and Methods

### Gene-based tests with multiple phenotypes for a single study

Suppose we intend to conduct a meta-analysis with *S* studies each having *K* phenotypes. For the *s^th^* study *n_s_* subjects are genotyped in a region that has *m_s_* variants. Let *y_ks_* = (*y*_1*ks*_,…, *y_n_s_ks_*)^*T*^ be the *n_s_*×1 vector for the *k^th^* phenotype on *n_s_* individuals in the *s^th^* study; *G_js_* = (*g*_1*js*_,…,*g_n_s_js_*)^*T*^ is an *n_s_*×1 vector of the minor allele counts (0, 1, or 2 variant alleles) for variant *j* and *G_s_* = (*G*_1*s*_,…,*G_m_s_s_*) is an *n_s_×m_s_* genotype matrix for the *m_s_* genetic variants in the target gene/region. For a gene-based multiple phenotype test, we consider the following regression model

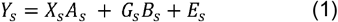

where *Y_s_* = *(y_1s_,…, y_Ks_)* is an *n_s_×K* phenotype matrix of *n_s_* individuals and *K* phenotypes; *B_s_* = (*β_jks_*) is an *m_s_×K* matrix where *β_jks_* is the regression coefficient of phenotype *k* on *G_js_*; *A_s_* is a *q_s_×K* matrix of regression coefficients for non-genetic covariates *X_s_*; and *E_s_* is an *n_s_×K* matrix of the error terms. The null hypothesis of no genetic association between variants in the region and the phenotype is H_0_: *β_jks_* = 0 for all *j* and *k*.

Let 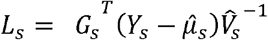 be the *m_s_* × *K* score matrix for the *s^th^* study where 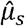 is an *n_s_×K* matrix of the estimated mean of *Y_s_* under the null hypothesis of no association, and 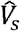 is the *K x K* estimated null residual covariance matrix among the *K* phenotypes in the *s^th^* study. To test the null hypothesis of no association, we use a variance component test. Under the mixed effect model set-up, we assume that the vectorized form of matrix *B_s_*, represented as *vec*(*B_s_*), follows a distribution with mean 0 and variance *τ*^2^*Σ_G_*⊗*Σ_P_* (for details on *Σ_G_* and *Σ_P_* see below), where ⊗ is a Kronecker product. The null hypothesis of no genetic association hence can be written as H_0_: *τ* = 0. The corresponding score-test statistic is

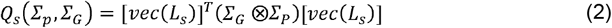

Under the null hypothesis, *vec*(*L_s_*) asymptotically follows *N*(0, *Φ_s_*) where *Φ_s_* is the phenotype-adjusted variant relationship matrix

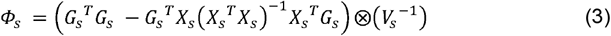

and *Q_s_*(*Σ_p_,Σ_G_*) asymptotically follows a mixture of chi-squares distribution. The mixing parameters are the eigenvalues of *RΦ_s_R^T^* where (*Σ_G_*⊗*Σ_P_*) = *RR^T^*.

The kernel *Σ_G_* represents the effect sizes of the variants to a phenotype. In general, *Σ_G_* is assumed to be a sandwich matrix *WR_G_W*, where *W* = *diag*(*w*_1_,…,*w_m_s__*)^*T*^ is a diagonal matrix for the variant-weighting, and 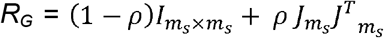 is a compound symmetric correlation matrix, where *I_m_s_×m_s__* is an *m_s_×m_s_* identity matrix and *J_m_s__* = (1,1,…,1)^*T*^ is an *m_s_*×1 vector with all elements being unity. This model can cover a wide range of scenarios of the genetic effect distribution. For example, with one phenotype (*K* = 1), if *ρ*=1 (i.e. *R_G_*=*J_m_s__J^T^m_s_*), which assumes homogeneous effects of the variants to the phenotype, the test reduces to a Burden test(Li & Leal, 2008; Madsen & Browning, 2009). Similarly if *ρ* = 0 (i.e. *R_G_* = *I_m_s_×m_s__*), the test is equivalent to a SKAT test(M. C. Wu et al., 2011).

The kernel *Σ_P_* represents the effect sizes of a variant on the phenotypes. For example, under the assumption that the genetic effects of a variant on each phenotype are independent, we can use *Σ_P_* = *I_K×K_*, which is numerically equivalent to Het-MAAUSS(Selyeong Lee et al., 2017). If the covariance structure in phenotypes is assumed to follow genetic effect residual covariance, we can use 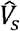 as *Σ_P_*, which results in the test equivalent to GAMuT(Broadaway et al., 2016), MSKAT(B. Wu & Pankow, 2016) and DKAT(Zhan et al., 2017).

### Input summary statistics from each study for meta-analysis

Single-variant meta-analyses are conducted with single-variant summary statistics, such as the estimated effect sizes and their standard errors. For region based tests with a single phenotype, Lee et al.(Seunggeun Lee, Teslovich, Boehnke, & Lin, 2013) showed that the score statistics of the variants, minor allele frequencies (MAFs) and the variant relationship matrix can be used as summary statistics for meta-analysis. With multiple phenotypes, the multivariate forms of these summary statistics from each study are needed. In particular from the s^th^ study, the score matrix, *L_s_*, the phenotype-adjusted variant relationship matrix, *Φ_s_*, the residual covariance structure of the phenotypes, 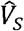, and the MAFs of the variants in the region are needed for meta-analysis.

### Meta-MultiSKAT: Meta-analysis of gene-based tests with multiple phenotypes

For simplicity, here we assume that all variants and phenotypes are observed in all *S* studies, so that *m* = *m_1_* =…= *m_s_*. We will relax this assumption later. Suppose summary statistics (*L_s_, Φ_s_*), *s* = 1,…, *S* are provided by *S* studies. We first construct the meta-score-vector as *L_meta_* = (*vec*(*L*_1_)^*T*^, *vec*(*L*_2_)^*T*^ … *vec*(*L_s_*)^*T*^)^*T*^. The variance component test statistic for meta-analysis is

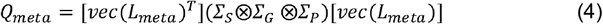

Under the null hypothesis, *Q_meta_* follows a mixture of chi-square and the corresponding p-value can be obtained by inverting the characteristic function (See Appendix A for details).

Here we have introduced another kernel *Σ_S_*. Similarly as the other kernels, *Σ_S_* models the heterogeneity between the effects of the contributing studies. In particular we will consider two special structures of *Σ_S_*:

*Homogeneous*, 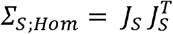, which assumes that across the *S* studies the effects of the variants on all the phenotypes are the same (homogeneous).

*Heterogeneous, Σ_S;Het_ = I_S_*, which assumes that across the *S* studies the effects of the variants on the phenotypes are uncorrelated or heterogeneous.

The test statistic (4) assumes that the kernels *Σ_G_* and *Σ_P_* are the same across studies. This assumption is restrictive since different studies might be analyzed with different hypotheses, reflected in different *Σ_G_, Σ_P_* across studies. This can be resolved by modifying (4) as

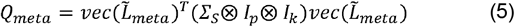

where 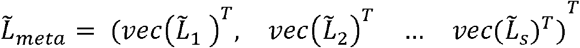 and 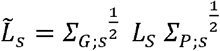 represents the kernelized scores incorporating study specific *Σ_G;S_* and *Σ_P;S_* for the *s^th^* study. (See Appendix B for details)

#### Variant weighting scheme

In region-based analysis, Wu et al.(MC et al., 2011) suggested MAF-based weighting scheme. To upweight the rare-variants, they proposed to use Beta(1,25) weights. When the homogeneity across the studies are assumed (i.e. *Σ_S_* = *Σ_S;Hom_*), pooled MAFs across studies can be used to generate weights for variants. For *Σ_S_* = *Σ_S;Het_*, we use study specific weights obtained using MAF’s of each study. Alternatively, functional scores, such as CADD(Kircher et al., 2014) and Eigen(Ionita-Laza, Capanu, De Rubeis, McCallum, & Buxbaum, 2014) can be used to upweight functionally important variants. In addition to using the MAF-based weighting, we have also explored the use of CADD scores as weights for variants in the meta-analysis of MGI and SardiNIA datasets.

### Combined effect of common and rare variants (Meta-MultiSKAT-Common-Rare)

The default setting for SKAT type tests (SKAT, MultiSKAT and Meta-MultiSKAT) is to use a MAF-based weighting scheme that up-weights the contribution of the rare variants and down-weights that of common variants. When there are common variants in the region associated with the phenotype, this weighting scheme can lead to a loss in power. Similar to Ionita-Laza et al.(Ionita-Laza, Lee, Makarov, Buxbaum, & Lin, 2013), we propose a test of the combined effects of common and rare variants on the phenotype. As in equation (4), the Meta-MultiSKAT test statistic is given by the quadratic form, *Q_meta_* = *vec*(*L_meta_*)^*T*^*K vec*(*L_meta_*), where *K* = (*Σ_S_*⊗*Σ_G_* ⊗*Σ_P_*). Given each study MAF, we compute the pooled MAF and using a cut-off on that we partition the variants into common and rare. In practice, cut-offs like 5% MAF or 1% MAF are commonly used. To explicitly separate the effects of common and rare variants, we construct the test statistic separately for common and rare variant. In particular, we construct

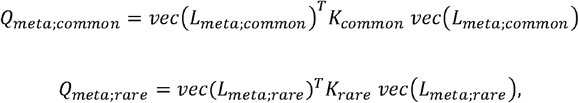

where *L_meta;common_* and *K_common_* (alternatively *L_meta;rare_* and *K_rare_*) are constructed using common variants (alternatively rare variants) only. The two matrices *K_common_* and *K_rare_* only differ in terms of the underlying *Σ_G_* matrices and we can allow different weighting schemes for the *Σ_G_* kernels corresponding to common and rare variants. In particular, here, we use Beta(0.5,0.5) weights for the common variants and Beta(1,25) weights for the common variants.

The combined sum (Meta-MultiSKAT-Common-Rare) is then constructed as

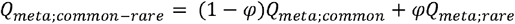

with a given weight *φ*. A simple approach, as used in Ionita-Laza et al.(Ionita-Laza et al., 2013), is to select *φ* such that the rare and common variants contribute equally to the test statistics, i.e. 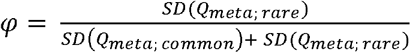. The asymptotic p-value of *Q_meta;common–rare_* can be calculated from a mixture of chi-squared distribution, similar to the previous discussion.

### Kinship adjustment within studies

Individual studies might require adjustment for kinship if there are related individuals within the study. For instance, if study s has related individuals with kinship matrix Ψ, co-heritability matrix *V_g;s_* and the shared non-genetic effect matrix *V_e;s_* then we construct scores as:

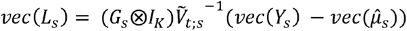

Where 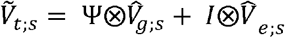. represents the estimated total covariance matrix for *Y_s_*.

### Discrepant phenotypes and genotypes across studies

The studies included in the meta-analysis may not have exactly the same set of variants genotyped (or sequenced). In particular, some variants may be observed in only a subset of studies. If variant *j* was not observed in study *s*, we set the (*j,k*)^*th*^ element in *L_s_* (k = 1, 2,…, K) and the corresponding elements in *Φ_s_* to be zero, which implies that the studies with missing data do not contribute to the score statistic. This also corresponds to imputing the missing data with the respective mean under the null hypothesis of no association. Using the same framework, if phenotype *k* in study s was not collected, we set the (*j,k*)^*th*^ element in *L_s_*(*j* = 1, 2,., *m_s_*) and the corresponding elements in *Φ_s_* to be zero. As above, this corresponds to the null hypothesis that the missing phenotype is not associated with the region of interest.

### Minimum p-value-based omnibus tests: Meta-Hom, Meta-Het, and Meta-Com

The Meta-MultiSKAT model and tests have three parameters *Σ_S_, Σ_P_* and *Σ_G_* that are absent in the null model. Since this is a score test, these parameters cannot be estimated from the data. One possible solution is to select them based on prior information, reflecting a specific hypothesis about the underlying model of association. However if the selected values do not reflect the true model, then the corresponding test might be underpowered(Selyeong Lee et al., 2017; Ray et al., 2016). To overcome such issues, minimum p-value-based omnibus tests have been proposed, which aggregate results across different values of the parameters to produce robust results(Dutta et al., 2019; Engel et al., 2013; He, Xu, Lee, & Ionita-Laza, 2017; Urrutia et al., 2015; Zhan et al., 2017). Here we use the same strategy to formulate robust tests across different choices of *Σ_S_, Σ_P_* and *Σ_G_*.

We first calculate p-values from different choices of (*Γ_S_, Σ_P_, Σ_G_*) and obtain the minimum of these p-values. Since the tests are correlated, using a Bonferroni correction can result in conservative type-I error and low power. Instead, we use a fast resampling approach to estimate the null correlation between the tests being aggregated and subsequently use a copula approach to estimate the p-value of the minimum p-value test statistic (See Appendix C for details)(Demarta & McNeil, 2007; Dutta et al., 2019). This approach has also been used previously to integrate information from multiple functional annotations(He et al., 2017). Specifically we consider the following tests:

Meta-Hom: minimum p-value of Meta-MultiSKAT tests with *Σ_S_* = *Σ_S;Hom_* across different choices of *Σ_P_* and *Σ_G_*. Specifically, we consider the following four different choices of (*Σ_P_, Σ_G_*)

- 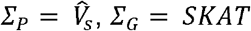
- 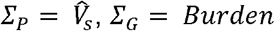
- *Σ_P_* = *I_K_, Σ_G_* = *SKAT*
- *Σ_P_* = *I_K_, Σ_G_* = *Burden*

The minimum p-value across these four tests will be used as the test statistic to evaluate the associations.

Meta-Het: minimum p-value of Meta-MultiSKAT tests with *Σ_S_* = *Σ_S;Het_* across different choices of *Σ_P_* and *Σ_G_*. We will use the same four sets of (*Σ_P_, Σ_G_*) as in Meta-Hom.

Meta-Com: combined test of Meta-Het and Meta-Het. We use the minimum p-value of the tests used in Meta-Hom and Meta-Het as test statistics.

## Simulations

We carried out extensive simulation studies to evaluate the type I error rate and power of Meta-MultiSKAT tests. For type-I error simulations and all power simulations, we generated 10,000 chromosomes over 1Mb-regions using a coalescent simulator with a European ancestry model(Schaffner, Stephen F et al., 2005). Because the average total exon length of a gene is about 3 kbps, we randomly selected a 3 kb region for each simulated dataset to test for associations.

### Simulation setting within individual study

In the *s^th^* study, we generate *K* phenotypes according to the linear model:

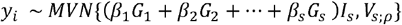

where *V_s;ρ_* is the covariance of the non-systematic error term. We use *V_s;ρ_* to define level of residual covariance between the traits. The matrix *V_s;ρ_* is set to be compound symmetric throughout all the simulation settings with varying values of the correlation parameter *ρ* (low correlation: ρ = 0.3; mild correlation: ρ = 0.5; high correlation: ρ = 0.7). *I_s_* is a *K*×1 indicator vector, which has 1 when the corresponding phenotype is associated with the region and 0 otherwise. Throughout our simulations we set *I_s_* = (1,1,1,0,0)′ meaning the first 3 phenotypes are associated with the region of interest in a particular study.

For estimating type-1 error rates we set *β_i_* = 0 for all the variants in all the studies. For power simulation, we used two different settings. In the first setting, to estimate the power of Meta-MultiSKAT as a rare-variant test, we set 30% of the rare variants (MAF ≤ 1%) to be causal. Next, to estimate the performance of Meta-MultiSKAT as a test of combined effects of common and rare variants, we set 30% of all variants (common or rare) in the region to be causal. We modeled rare variants to have stronger association with the phenotypes than the common variants by setting |*β_j_*| = *c*log_10_|*MAF_j_*| with *c* = 0.2 for all the simulation scenarios. For both the settings, as mentioned earlier, the first three among the five phenotypes in each study were associated with the region of interest.

### Simulation settings across studies

Throughout our simulations we have used settings which consist of three studies on European samples with five phenotypes of interest. The sample sizes for the studies were 2000, 2000 and 1000 respectively. To assess the performance of Meta-MultiSKAT under scenarios of missing data, we considered the following 3 scenarios:

Scenario A: all the individuals in each of the study have complete information on 5 correlated phenotypes

Scenario B: 10% samples (chosen completely at random) in the 3^rd^ study have information on 4 phenotypes only. This means, 100 samples in study 3 have information on 4 phenotypes and misses information on 1 phenotype, while the rest 900 samples have information on all the 5 phenotypes. All the 2000 samples in study 1 and 2 have complete information on all the 5 phenotypes.

Scenario C: The 5^th^ phenotype for study 3 is missing for all the samples. For study 1 and 2, all the 2000 samples have complete information on all the 5 phenotypes.

For these above scenarios, in addition to the Meta-MultiSKAT tests (Meta-Hom, Meta-Het and Meta-Com), we evaluated the following single phenotype-based approaches:

1. MinPhen-Het: Bonferroni-adjusted minimum p-value from the single phenotype region-based meta-analysis using Heterogeneous Meta-SKAT-O (Het-Meta-SKAT-O)
2. MinPhen-Hom: Bonferroni-adjusted minimum p-value from the single phenotype region-based meta-analysis using Homogeneous Meta-SKAT-O (Hom-Meta-SKAT-O)

### Meta-analysis of white blood cell traits

To investigate the pleiotropic roles of low frequency and rare-variants on WBC subtypes, we analyzed data collected under the Michigan Genomics Initiative (MGI study)(Fritsche et al., 2018) Phase 2 (data-freeze on December 2017) and the SardiNIA(Vacca et al., 2006) study. Data on four WBC subtypes percentages were included in the analysis: lymphocyte, monocyte, basophil and eosinophil. We excluded the data on percentage of neutrophils since it was highly correlated with lymphocytes (absolute value of correlation > 0.9 in both MGI and SardiNIA). European samples with at most two phenotypes missing were included in the analysis for each of the studies. In all, we included 11,049 and 5,899 samples from the MGI and the SardiNIA studies, respectively (Table 1).

**Table 1:** Sample sizes for each phenotype in each study for MGI-SardiNIA meta-analysis.

| Phenotype | N (MGI) | N (SardiNIA) | N (Total) |
| --- | --- | --- | --- |
| Lymphocyte % | 11049 | 5895 | 16944 |
| Monocyte % | 11049 | 5876 | 16925 |
| Basophil % | 11038 | 5866 | 16904 |
| Eosinophil % | 11037 | 5795 | 16830 |
| Complete Cases | 11035 | 5735 | 16770 |
| Total Samples | 11049 | 5898 | 16948 |

We annotated protein-coding variants and a region of 20kb (± 10kb) around them to genes using Variant Effect Predictor(McLaren et al., 2016) software.

Within each study, we included age, sex, and study specific top four principal components (PC) as fixed effect covariates in the analysis. In each study each of the four WBC subtypes were adjusted for the corresponding covariates and the residuals were quantile-normalized. Further, we estimated the kinship between the subjects in each study using KING(Manichaikul et al., 2010) and estimated the co-heritability matrix of the phenotypes using PHENIX(Dahl et al., 2016). The inverse normalized residuals were then used in region-based multiple phenotype analysis (Multi-SKAT with kinship correction). The required summary statistics were calculated from the individual tests.

We conducted three sets of analysis with the extracted summary statistics. First to test the rare-variant associations of the phenotypes, we used Meta-MultiSKAT tests (Meta-Het, Met-Hom and Meta-Com) to test groups of protein-coding variants with pooled MAF ≤ 1%. We only included the groups that had at least three variants and a total minor allele count of 5. We used a Beta(1,25) weighting scheme to upweight the effect of the rare variants. Next, to test the combined effect of common and rare variants, we used the Meta-MultiSKAT-Common-Rare versions of the above tests with groups of protein-coding variants without any MAF cutoff. This means both common (MAF > 1%) and rare variants (MAF ≤ 1%) were present in the regions tested. For the rare variants we used Beta(1,25) weights and for the common variants we used Beta(0.5,0.5) weights (see Methods). Further, we annotated CADD scores for all the variants (common and rare) using ANNOVAR(Wang, Li, & Hakonarson, 2010). We used these scores as weights in the genotype kernel *Σ_G_* and performed the above Meta-MultiSKAT tests.

## Results

### Type-I error

For type-I error simulations, we simulated 10^7^ independent datasets with three studies each having five phenotypes with a compound symmetric null residual covariance structure with off-diagonal elements being equal to 0.5, i.e. *V*_*s*;0.5_. The MAF spectrum for the population allele frequencies shows that the majority of the simulated variants are rare (MAF ≤ 1%). We estimated the type-1 error rate as the proportion of p-values less than the specified α levels, with α set at 10^−4^, 10^−5^ and 2.5×10^−6^.

Type-I error rates were well maintained at all α levels. For example, at α = 2.5×10^−6^, the largest estimated type-I error rate for any of the Meta-MultiSKAT tests was 2.7×10^−6^, which was well within the estimated 95% confidence interval (Table 2).

**Table 2:** Estimated Type-1 error rates for Meta-MultiSKAT tests.

| $\alpha$ | Meta-Hom | Meta-Het | Meta-Com |
| --- | --- | --- | --- |
| $2.5 \times 10^{-6}$ | $2.3 \times 10^{-6}$ | $2.6 \times 10^{-6}$ | $2.7 \times 10^{-6}$ |
| $1 \times 10^{-5}$ | $1.1 \times 10^{-5}$ | $1.1 \times 10^{-5}$ | $1.2 \times 10^{-5}$ |
| $1 \times 10^{-4}$ | $1.2 \times 10^{-4}$ | $1.2 \times 10^{-4}$ | $1.3 \times 10^{-4}$ |

### Power

We compared the empirical power of Meta-MultiSKAT tests with two possible existing approaches: minimum of the single phenotype MetaSKAT p-values (MinPhen-Hom and MinPhen-Het). For each simulation setting, we generated 1000 datasets and estimated the empirical power as the proportion of p-values less than 2.5×10^−6^, reflecting the Bonferroni correction for testing 20,000 independent genes.

In power simulations, the first scenario considered the case that each study has the same set of causal variants and all of them are trait-increasing. Meta-Hom and Meta-Com had the highest powers in all scenarios while the power for Meta-Het is lower (Figure 1). Also, there was a slight overall decrease in power from scenario A through scenario C. We expect this decrease in power since there is an increase of the amount of missing-ness in the scenarios A through C, though the power decrease is small (maximum relative decrease in empirical power < 1%). Overall power of all the methods was higher when the correlation is high (ρ = 0.7).

**Figure 1:**
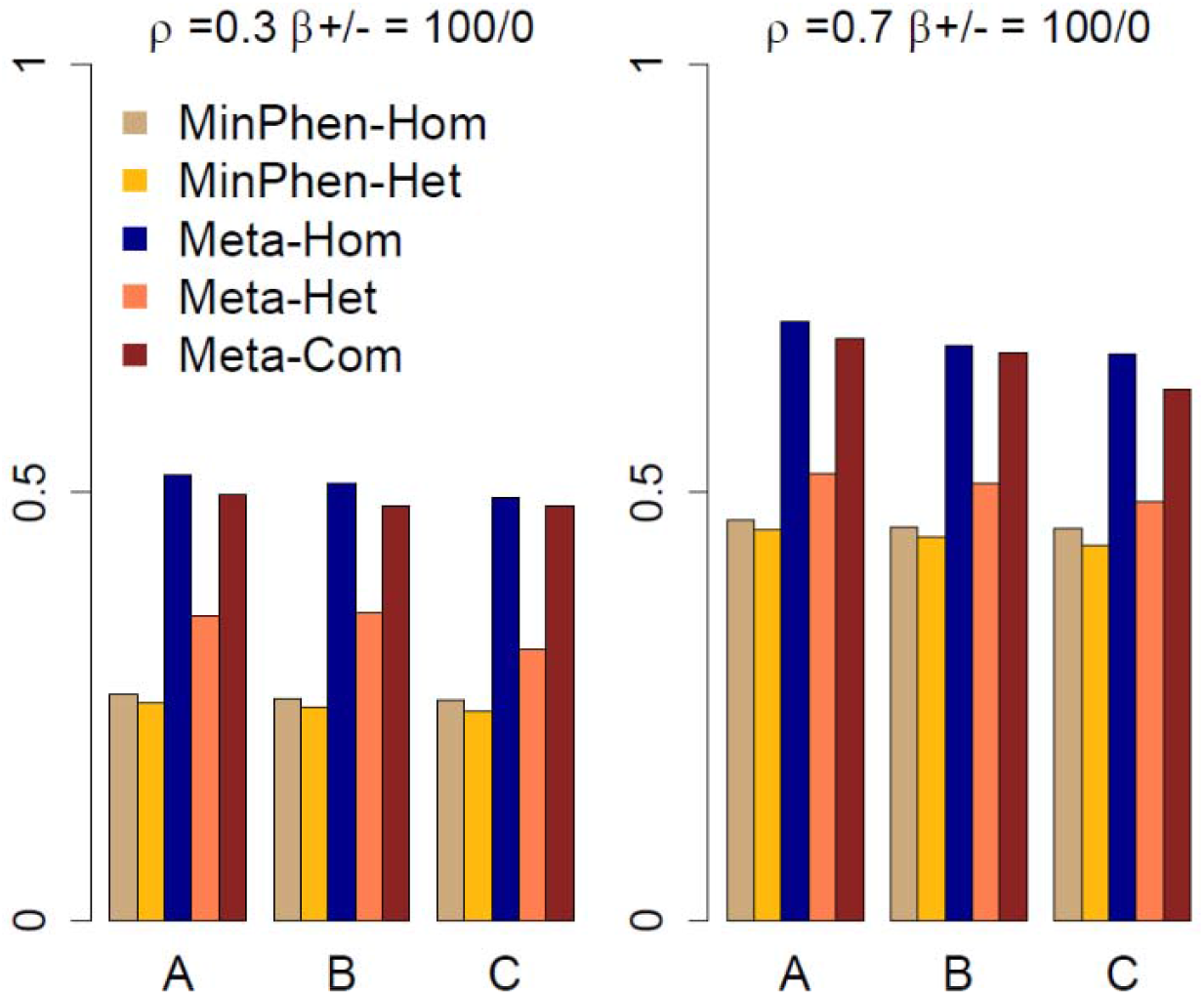
Power for Meta-MultiSKAT tests compared with the existing methods when the set of causal variants is the same across different studies and has the same direction of effect. Empirical power for Meta-Hom, Meta-Het and Meta-Com plotted for 3 different scenarios compared against MinPhen-Hom and MinPhen-Het (See Simulations for details). Left panel shows the results for low correlation (ρ = 0.3) among the phenotypes and right panel shows the results for high correlation (ρ = 0.7).

Next, we considered a heterogeneous situation in which causal variants for each study were randomly selected so only small percentage of causal variants were shared among studies (Figure 2). As expected, Meta-Het and Meta-Com had high power among the tests being compared. Meta-Hom was underpowered compared to these tests, while MinPhen-Hom and MinPhen-Het had lower power than the rest.

**Figure 2:**
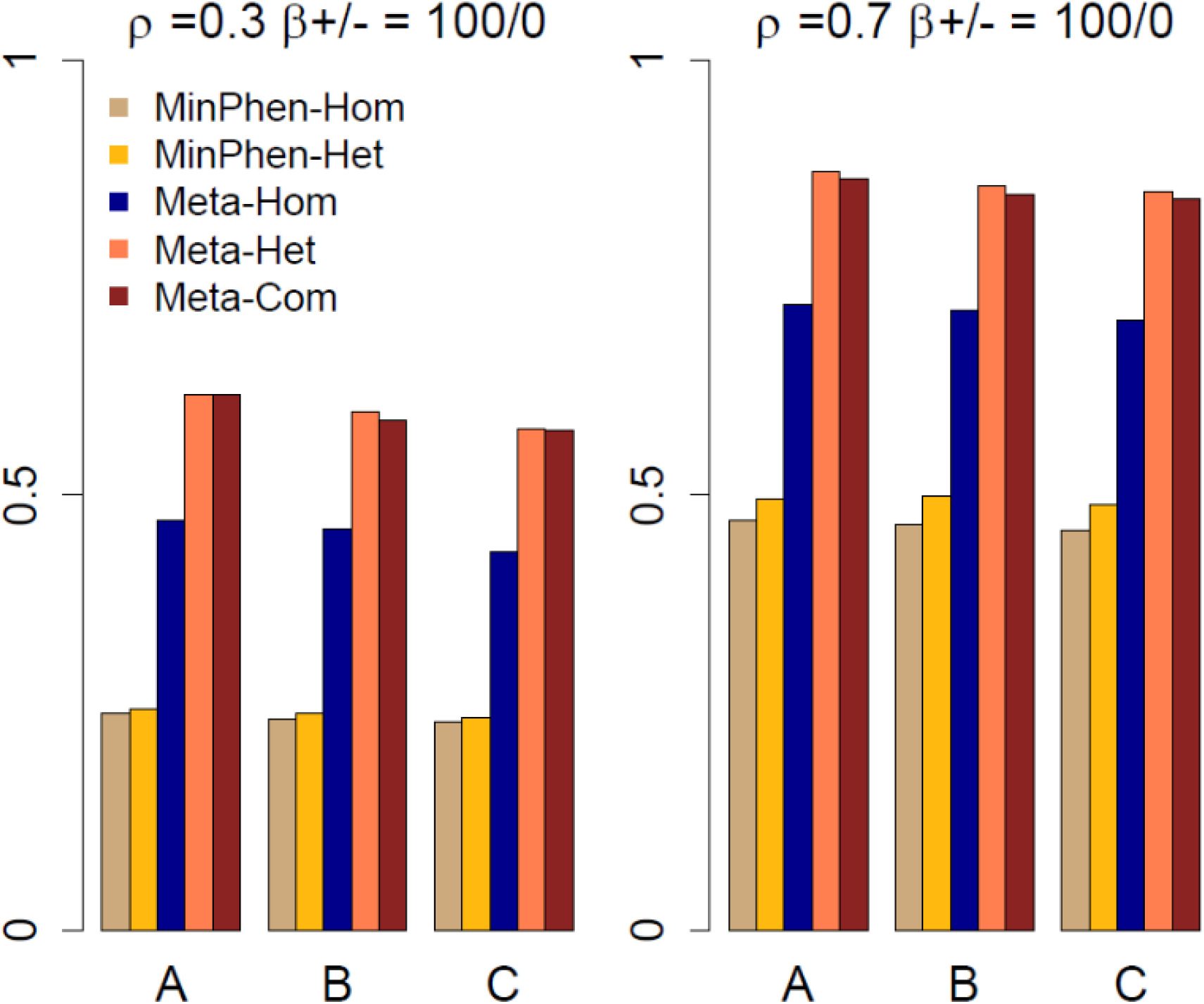
Power for Meta-MultiSKAT tests compared with the existing methods when the set of causal variants is randomly chosen for each study and has the same direction of effect. Empirical power for Meta-Hom, Meta-Het and Meta-Com plotted for 3 different scenarios compared against MinPhen-Hom and MinPhen-Het (See Simulations for details). Left panel shows the results for low correlation (ρ = 0.3) among the phenotypes and right panel shows the results for high correlation (ρ = 0.7).

We then assumed that the causal variants for each study are chosen randomly within the region and 20% of the variants are trait-decreasing (80% are trait increasing) (Figure 3). Similar to the previous scenario, Meta-Het and Meta-Com had higher power than the rest of the tests. MinPhen-Hom and MinPhen-Het had lower power of detecting association signals, and Meta-Hom consistently had the lowest power across all the settings.

**Figure 3:**
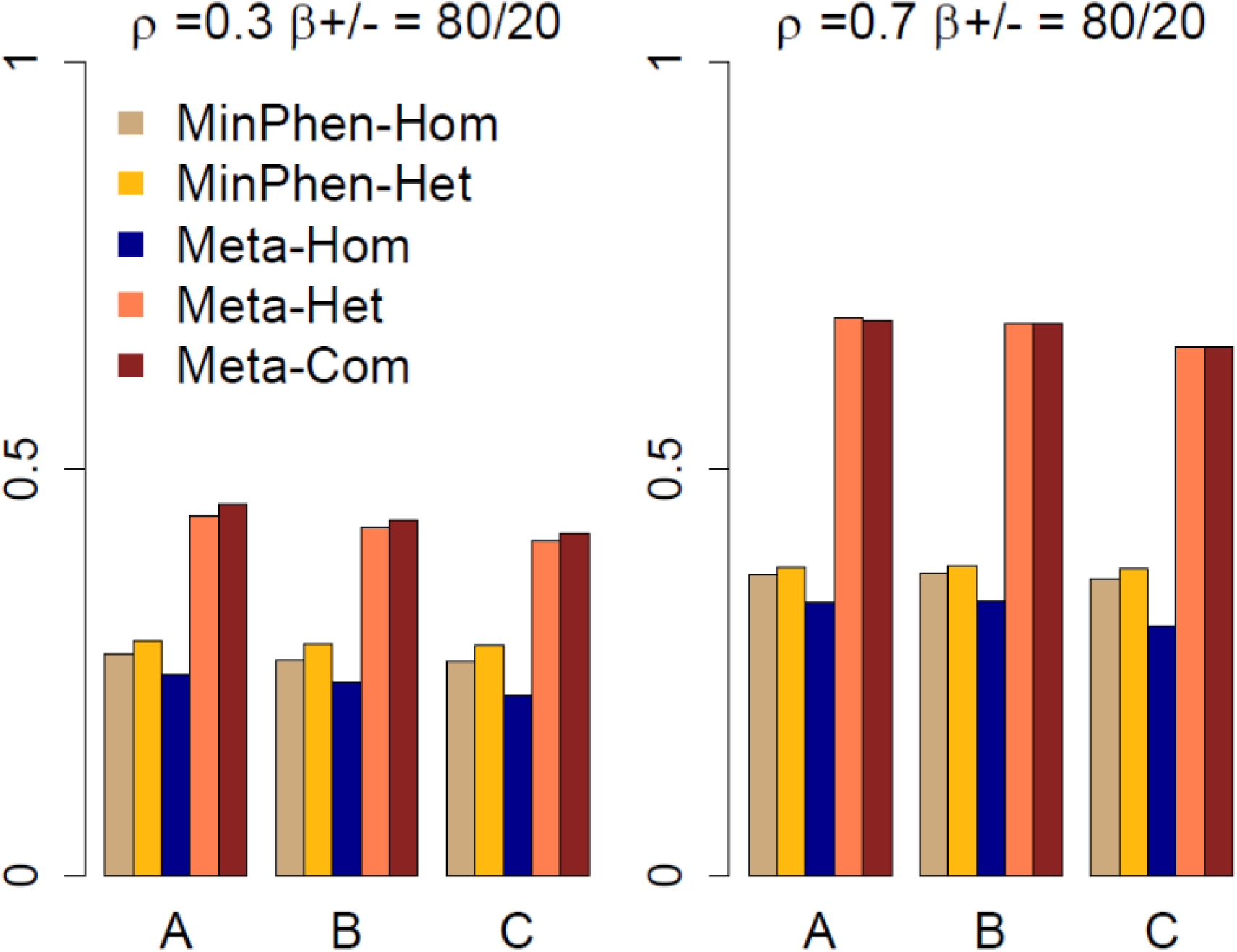
Power for Meta-MultiSKAT tests compared with the existing methods when the set of causal variants is randomly chosen for each study and 20% of the causal variants are trait-decreasing. Empirical power for Meta-Hom, Meta-Het and Meta-Com plotted for 3 different scenarios compared against MinPhen-Hom and MinPhen-Het (See Simulations for details). Left panel shows the results for low correlation (ρ = 0.3) among the phenotypes and right panel shows the results for high correlation (ρ = 0.7).

Next we considered a situation where the correlation structure among the phenotypes across studies varies. For the 1^st^ and 2^nd^ study the correlation among the 5 phenotypes is high (ρ = 0.7) while for the 3^rd^ study, the correlation among the 5 phenotypes is moderate (ρ = 0.5). Similar to the previous cases, Meta-Het and Meta-Com maintained higher power than the rest of the tests (Figure 4). As before, Meta-Hom performed poorly when 20% of the causal variants are trait-decreasing.

**Figure 4:**
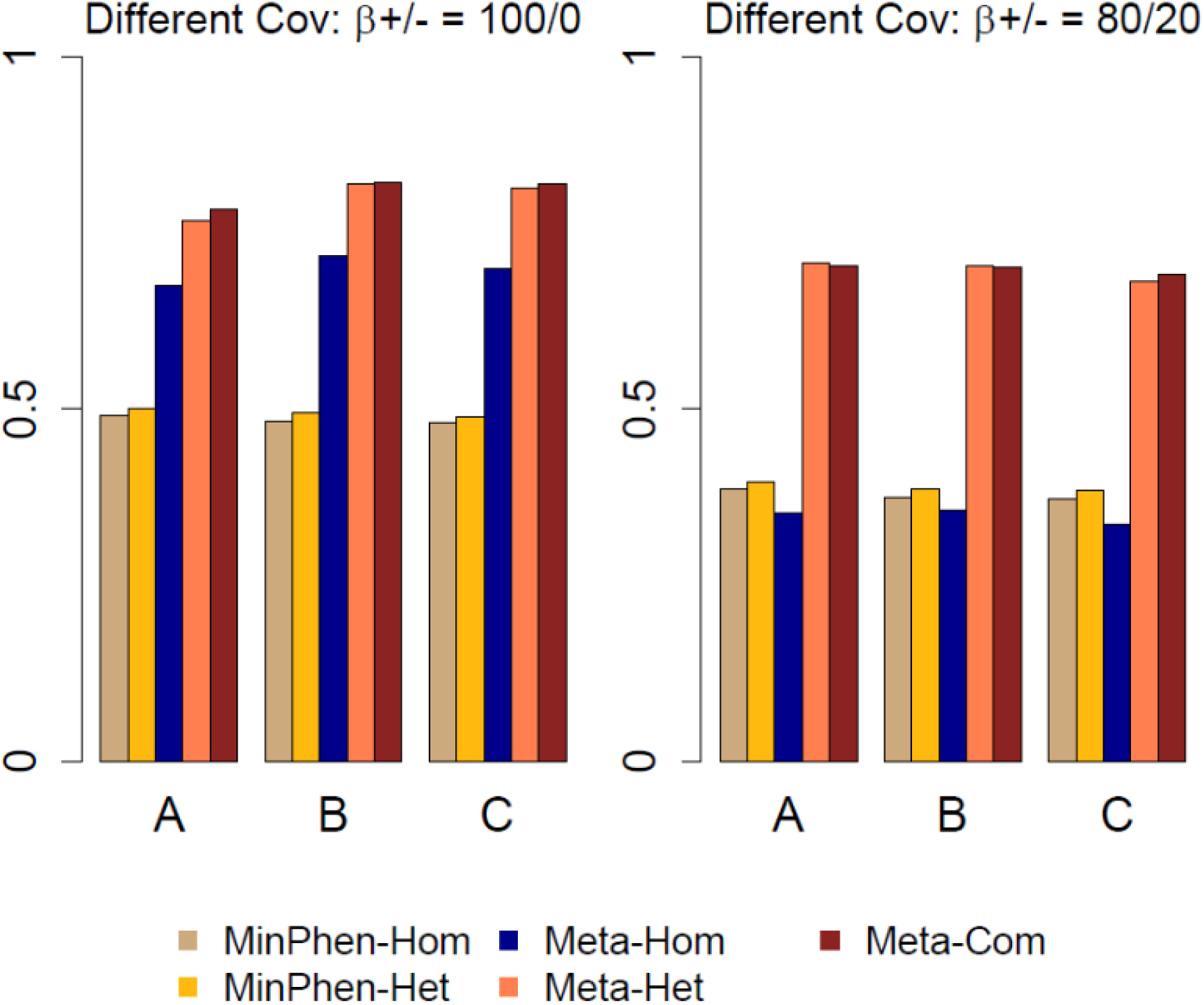
Power for Meta-MultiSKAT tests compared with the existing methods when the set of causal variants is randomly chosen for each study and the studies have different covariance structure across the phenotypes. Empirical power for Meta-Hom, Meta-Het and Meta-Com plotted for 3 different scenarios compared against MinPhen-Hom and MinPhen-Het (See Simulations for details). Left panel shows the results when all the causal variants are trait increasing; right panel shows the same when 20% of the causal variants are trait-decreasing.

We further estimated type-1 error and power for the Meta-MultiSKAT-Common-Rare versions of the tests. The results are shown in Supplemental Table S3, Supplemental Figure S2 and S3. Type-I error was well maintained at different levels and the patterns of estimated power remained the same.

Overall our simulations show that Meta-MultiSKAT tests can improve power over the existing single phenotype-based meta-analysis approaches, while controlling type-I error rates. In particular, Meta-Com maintains robust power across all the scenarios regardless of the underlying genetic model.

### Meta-analysis of WBC subtype traits

White blood cells (WBCs) are major cellular components of the human immune system. They have been found to be associated with risk of cardiovascular disease(J. H. Kim, Lim, Park, Jang, & Choi, 2017) and cancer mortality(Erlinger, Muntner, & Helzlsouer, 2004) among others. Certain disease risk factors including high blood pressure, cigarette smoking, adiposity and increased levels of plasma inflammatory markers have been reported to be linked to elevated WBC counts(Hasegawa, Negishi, & Deguchi, 2002; Mu oz et al., 2012). WBCs are classified into subtypes according to the functionality and morphology. Abundances (counts or percentage) of these WBC subtypes have been found to be important biomarkers for diseases including COPD(D. K. Kim et al., 2012) and rheumatoid arthritis(Salomon et al., 2017), and several GWAS have identified genetic variants associated with them(Astle et al., 2016; Crosslin et al., 2013; Kanai et al., 2018; Keller et al., 2014). In this analysis, we tested the abundances of lymphocyte, monocyte, basophil and eosinophil. Correlations among the phenotypes are shown in Supplemental Figure S4. There are more strong correlations in MGI samples.

We applied Meta-MultiSKAT tests to the analysis of WBC subtypes from the MGI and SardiNIA studies (See Methods for details). In particular, we applied Meta-Het, Meta-Hom and Meta-Com tests along with MinPhen-Hom and MinPhen-Het. We also evaluated the single-phenotype tests and multiple-phenotype tests (Multi-SKAT) for each study (Supplemental Table S1).

#### Results for rare variants with MAF-based weighting

First, we used the MAF-based weighting scheme to upweight the rare variants as suggested by Wu et al(MC et al., 2011). Using the variants with pooled MAF ≤ 1%, we used Beta(1,25) weights. Overall 5,109 genes with at least 3 variants and a total minor allele count > 5 were tested. This produces a Bonferroni cut-off of 9.8×10^−6^ (approximately 1×10^−5^)

The QQ-plots shown in Figure 5 corresponding to the Meta-MultiSKAT tests do not show any indication of inflation (genomic control varying from 0.998 to 1.003). Table 3 shows the genes that had p-values less than 1×10^−5^ for at least one of the tests. Two genes *PRG2* (p-value = 5.9×10^−7^) and *RP11-872D17.8* (p-value = 1.7×10^−7^) were identified as significant by Meta-MultiSKAT tests while the p-values for the existing tests did not reach significance.

**Figure 5:**
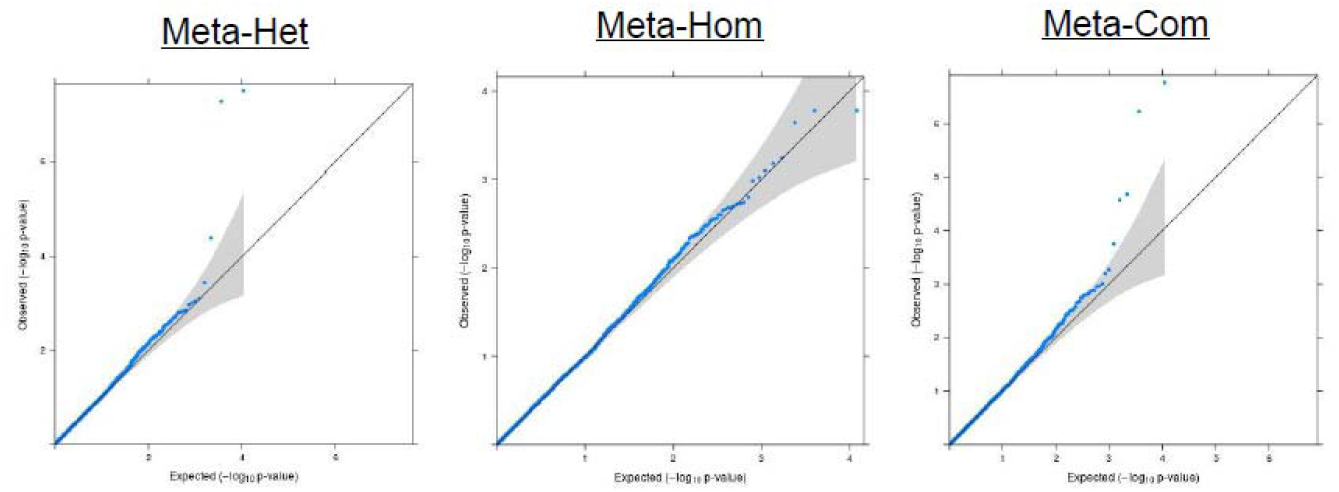
QQ plots for the Meta-MultiSKAT (Meta-Het, Meta-Hom and Meta-Com respectively) p-values obtained from MGI-SardiNIA meta-analysis.

**Table 3:**
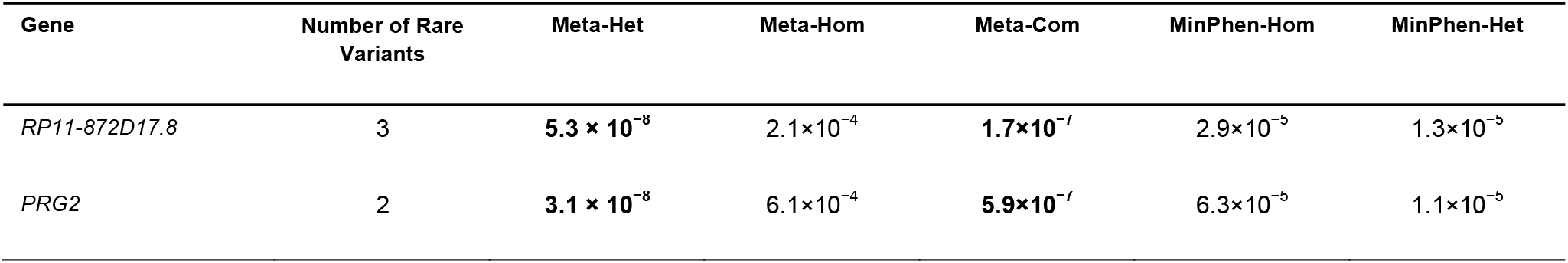
Genes/ regions identified by either of the Meta-MultiSKAT methods (Meta-Hom, Meta-Het or Meta-Com) or the existing approaches (MinPhen-Hom or MinPhen-Het). The p-values < 10^−5^ were marked in bold. Variants with pooled MAF ≤ 1% are included as rare.

| Gene | Number of Rare Variants | Meta-Het | Meta-Hom | Meta-Com | MinPhen-Hom | MinPhen-Het |
| --- | --- | --- | --- | --- | --- | --- |
| *RP11-872D17.8* | 3 | **$5.3 \times 10^{-8}$** | $2.1 \times 10^{-4}$ | **$1.7 \times 10^{-7}$** | $2.9 \times 10^{-5}$ | $1.3 \times 10^{-5}$ |
| *PRG2* | 2 | **$3.1 \times 10^{-8}$** | $6.1 \times 10^{-4}$ | **$5.9 \times 10^{-7}$** | $6.3 \times 10^{-5}$ | $1.1 \times 10^{-5}$ |

*PRG2* gene [MIM: 605601] encodes a protein, which is a major contributor to the crystalline core of the eosinophil granule. Multiple phenotype analysis (Multi-SKAT) shows evidence for a strong association in the SardiNIA study (p-value = 2.8×10^−7^) whereas the p-value in the MGI study (p-value = 0.76) does not show evidence for association (Supplemental Table S1). This signal is driven by the association of the gene with eosinophils in SardiNIA (SKAT-O p-value = 3.7 ×10^−7^; Supplemental Table S1). A low-frequency SNP at 11:57156106 (A/G; MAF 3% in SardiNIA), which is significantly associated with the eosinophil percentages (p-value = 9.3×10^−12^). This variant is only observed for the individuals in SardiNIA study and was not observed in MGI. The signal for *RP11-872D17.8*, an adjacent gene to *PRG2*, is also driven by the same variant.

Our results suggest that Meta-MultiSKAT can outperform the standard tests in detecting rare-variant associations while maintaining calibrated type-1 error rate. The genes *PRG2* and *RP11-872D17.8* were identified in a particular study and were again identified through Meta-MultiSKAT tests.

#### Results with combined effects of common and rare variant

To illustrate how Meta-MultiSKAT can test the combined effect of common and rare variants, we used Meta-MultiSKAT-Common-Rare version of each test to analyze the WBC data from the MGI and SardiNIA studies. All the variants, common and rare, were used in this analysis. The same two genes, *PRG2* and *RP11-872D17.8*, had p-values less than 1×10^−5^, but with different p-values along with *CCL24* and *IRF8* (Table 4).

**Table 4:** Genes/ regions identified by either of the Meta-MultiSKAT-Common-Rare methods (Meta-Hom, Meta-Het or Meta-Com). The p-values < 10^−5^ were marked in bold. Variants with pooled MAF ≤ 1% (> 1%) are included as rare (common).

| Gene | Number of<br>Common variants | Number of<br>Rare variants | Meta-Het | Meta-Hom | Meta-Com | MinPhen-Hom | MinPhen-Het |
| --- | --- | --- | --- | --- | --- | --- | --- |
| *IRF8* | 28 | 3 | $1.3 \times 10^{-4}$ | **$1.8 \times 10^{-10}$** | **$2.9 \times 10^{-10}$** | $7.4 \times 10^{-3}$ | $4.9 \times 10^{-4}$ |
| *PRG2* | 5 | 2 | **$5.7 \times 10^{-8}$** | $5.3 \times 10^{-5}$ | **$4.2 \times 10^{-8}$** | $6.3 \times 10^{-5}$ | $1.1 \times 10^{-5}$ |
| *CCL24* | 12 | 1 | **$4.6 \times 10^{-7}$** | $8.3 \times 10^{-4}$ | **$8.1 \times 10^{-7}$** | $8.2 \times 10^{-2}$ | $2.2 \times 10^{-4}$ |
| *RP11-872D17.8* | 9 | 3 | **$1.1 \times 10^{-6}$** | $1.1 \times 10^{-3}$ | **$2.4 \times 10^{-6}$** | $2.9 \times 10^{-5}$ | $1.3 \times 10^{-5}$ |

Among the genes that showed signal, Chemokine (C-C motif) ligand 24 gene (*CCL24* [MIM: 602495]) encodes a protein that interacts with chemokine receptor CCR3 to induce chemotaxis in eosinophils(White et al., 1997). This chemokine is also strongly chemotactic for resting T lymphocytes and slightly chemotactic for neutrophils(Salcedo et al., 2002). Multiple phenotype tests of *CCL24* did not reach significance in any of the individual studies (p-value in SardiNIA = 1.3×10^−2^; p-value in MGI = 2.0×10^−4^; Supplemental Table S1). But Meta-Het (p-value = 4.8×10^−7^) and Meta-Com (p-value = 8.1 ×10^−7^) are significant indicating the utility of meta-analysis to identify this signal.

Interferon regulatory factor 8 (*IRF8* [MIM: 601565]) at 16q24.1 has been previously reported as associated with several WBC subtype traits. *IRF8* has been found to be associated with monocyte count(Sichien et al., 2016) and has also been identified as a multiple sclerosis susceptibility locus(De Jager et al., 2009). Animal model studies showed that *IRF8* as a transcription factor plays an essential role in the regulation of lineage commitment during monocyte differentiation(Kurotaki et al., 2018; Yáñez, Ng, Hassanzadeh-Kiabi, & Goodridge, 2015). Astle et al(Astle et al., 2016) (2016) found several associations of *IRF8* with WBC subtype traits like Neutrophils (high correlation with Lymphocytes) and combinations (sum of neutrophil and basophil counts) in the UK Biobank. Meta-Hom (p-value= 1.8×10^−10^) and Meta-Com (p-value = 2.9×10^−10^) show evidence for strong association.

For *IRF8* (p-value = 2.9×10^−10^), *CCL24* (p-value = 8.1×10^−7^) and *PRG2* (p-value = 4.2×10^−8^) the combined effect of common and rare variants produces substantially more significant results compared to the MAF-based weighting with rare variants, without evidence of inflated false discoveries. In comparison, the p-values for *RP11-872D17.8* (p-value = 2.4×10^−6^) remain approximately of the same order of significance. The results from this analysis demonstrate that Meta-MultiSKAT can be applied as a region-based test for testing the combined effect of common and rare variants.

#### Results with CADD-score weighting with both common and rare variants

We reanalyzed the WBC data from the MGI and SardiNIA studies by weighting the variants with a functional score. Both common and rare variants were used in this analysis. We used CADD-scores as weights in *Σ_G_* = *WR_G_W*. The results are shown in Table 5. The same set of 4 genes identified using MAF-based weighting remained significant (p-value< 1×10^−5^), but with slightly different p-values. For *PRG2* (p-value = 1.2×10^−8^), *IRF8* (p-value = 1.6×10^−7^) and *CCL24* (p-value = 1.1×10^−6^), weighting by functional scores resulted in a slightly smaller p-value as compared to MAF-based weighting. For *RP11-872D17.8* (p-value = 2.4×10^−6^), however, the p-value remained nearly the same.

**Table 5:** Genes/ regions identified by either of the Meta-MultiSKAT methods (Meta-Hom, Meta-Het or Meta-Com) with functional (CADD) score weights were used for the variants. The p-values < 10^−5^ were marked in bold. Variants with pooled MAF ≤ 1% (> 1%) are included as rare (common).

| Gene | Number of<br>Common variants | Number of<br>Rare variants | Meta-Het | Meta-Hom | Meta-Com | MinPhen-Hom | MinPhen-Het |
| --- | --- | --- | --- | --- | --- | --- | --- |
| *IRF8* | 28 | 3 | $1.2 \times 10^{-5}$ | **$9.4 \times 10^{-8}$** | **$1.6 \times 10^{-7}$** | $7.4 \times 10^{-3}$ | $4.9 \times 10^{-4}$ |
| *PRG2* | 5 | 2 | **$7.4 \times 10^{-9}$** | $3.6 \times 10^{-4}$ | **$1.2 \times 10^{-8}$** | $6.3 \times 10^{-5}$ | $1.1 \times 10^{-5}$ |
| *CCL24* | 12 | 1 | **$8.2 \times 10^{-7}$** | $5.7 \times 10^{-4}$ | **$1.1 \times 10^{-6}$** | $8.2 \times 10^{-2}$ | $2.2 \times 10^{-4}$ |
| *RP11-872D17.8* | 9 | 3 | **$1.0 \times 10^{-6}$** | $1.1 \times 10^{-3}$ | **$2.3 \times 10^{-6}$** | $2.9 \times 10^{-5}$ | $1.3 \times 10^{-5}$ |

Further, Supplemental Table S1 lists the single phenotype and multiple phenotype p-values for these four genes in each study. It is to be noted that, no other gene had a p-value less than 1×10^−5^ in any of the single phenotype or multi-phenotype tests in each study.

As a further illustration, we performed Meta-MultiSKAT tests by masking lymphocyte data in SardiNIA and treating that as a missing phenotype (See Appendix D for details). The results (Supplemental Table S2) show that Meta-MultiSKAT has a robust power under such scenarios while controlling type-1 error.

### Computation Time

We estimated the computation time of Meta-MultiSKAT tests using simulated datasets on 3 studies (as described in the Simulations section) with 5 phenotypes and 50 genetic variants. We set the number of perturbation iterations to 1000. On average, Meta-Hom and Meta-Het tests required approximately 8 CPU-seconds (Intel Xeon 2.80 GHz) and Meta-Com required 12 CPU-sec. Analyzing the MGI and SardiNIA datasets, using the extracted summary statistics from each study, required about thirty CPU-hours when parallelized to 10 processes.

## Discussion

We propose a new method, Meta-MultiSKAT, which meta-analyzes region-based association of multiple phenotypes across studies. The model is based on study-specific summary statistics for the region and is flexible to accommodate a range of heterogeneity of genetic effects across studies. The simulation and the real data analysis results involving the summary statistics from MGI and SardiNIA demonstrate that Meta-MultiSKAT can substantially increase power compared to the existing tests and can identify additional association signals, while maintaining the desired type-I error rate. The method is implemented as an R-package (MetaMultiSKAT, see Web Resources).

We note that the test statistics, assuming homogeneous genetic effects, are essentially identical to joint analysis test statistics using all individual level data and accounting for study-specific covariate effects, resulting in nearly identical power using meta-analysis and joint analysis. Our power-simulations confirm this finding (Supplemental Figure S1).

For Meta-MultiSKAT tests with a given choice of *Σ_S_, Σ_P_* and *Σ_G_*, asymptotic p-values can be calculated. For Meta-Hom and Meta-Het, we are aggregating four such Meta-MultiSKAT tests for a given choice of *Σ_S_, Σ_P_* and *Σ_G_*. Although the corresponding p-values depend on a resampling scheme, they still can be calculated using a small number of perturbations. Similarly, p-values for Meta-Com that aggregates 8 Meta-MultiSKAT tests with a given choice of *Σ_S_, Σ_P_* and *Σ_G_* can also be calculated using a small number of resampling iterations. The reported computation times show that Meta-MultiSKAT tests are computationally manageable at a genome-wide level.

In addition, Meta-MultiSKAT retains the desirable properties of Multi-SKAT. For instance, Multi-SKAT effectively incorporates kinship information through a regression framework, allowing the use of the whole sample rather than only unrelated individuals in a particular study. Meta-MultiSKAT can use the kinship adjusted summary statistics from the Multi-SKAT tests across several studies to produce a test of association, in which kinship information for each study has been incorporated. This integration allows for the use of all samples for each of the studies, further augmenting statistical power.

The asymptotic p-value calculations for Meta-MultiSKAT rely on the normality assumption of the score vectors. When at least one pair of the phenotypes is very strongly correlated (i.e. absolute correlation > 0.9), this assumption may be violated. Currently, we do not have a mechanism to adaptively select an active set of phenotypes which might produce the optimal association signal for a particular region. Hence, we recommend that the data be pre-pruned for correlation and such strongly correlated phenotypes be excluded before analysis.

Currently, the framework of Meta-MultiSKAT is developed for continuous phenotypes. A direction of future research is to extend this framework for phenotypes that are a mixture of continuous and discrete types.

In summary, we have developed Meta-MultiSKAT, a meta-analysis method for testing rare-variant associations of multiple correlated phenotypes. Meta-MultiSKAT has robust power and can handle practical problems such as missing data and different covariance structures. The method provides a scalable and practical solution to test multiple phenotypes jointly and thus can contribute to detecting regions in the genome with pleiotropic effects.

## Acknowledgements

We would like to acknowledge Michigan Genomics Initiative and the SardiNIA studies for their help in providing the data. This study was supported by grants R01-HG008773 and R01-LM012535 (D.D. and S.L.). This research was also supported in part by the Intramural Research Program of the National Institute on Aging, National Institutes of Health (NIH) (contracts N01-AG-1-2109 and HHSN271201100005C), and by the National Institutes of Health (NHLBI grant HL117626).

## Web Resources

Online Mendelian Inheritance in Man (OMIM): http://www.omim.org

GitHub R-package: https://github.com/diptavo/MetaMultiSKAT

Variant Effect Predictor: http://grch37.ensembl.org/Homo_sapiens/Tools/VEP.

ANNOVAR: http://www.openbioinformatics.org/annovar/annovar_download.html

## Appendix

## Section A: P-value for Meta-MultiSKAT tests

In equation (4), under null hypothesis, *L_meta_* asymptotically follows *N*(0, *Φ_meta_*) where 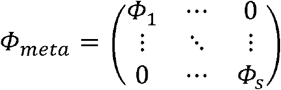. Hence, under the null hypothesis, *Q_meta_*, as in equation (4) follows a mixture of chi-squares. The mixing parameters of the distribution are eigenvalues of 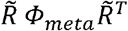 with 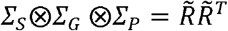. The p-value for this test can be obtained by inverting the characteristic function of the null distribution.

## Section B: Kernelized Scores

It is to be noted that equation (4) assumes that *Σ_G_, Σ_P_* are the same for individual studies. This assumption is restrictive as different studies might be analysed with different hypotheses, reflected in different *Σ_G_, Σ_P_* across studies. We can relax this assumption by using kernelized score matrix for each study in place of the score matrix. We construct the kernelized score statistic in each study as:

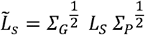

Under the null hypothesis of no association, 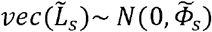 where 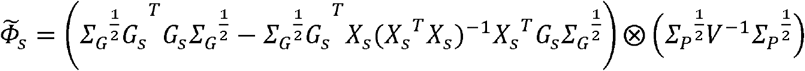 is the kernelized phenotype adjusted variant relationship matrix. Given 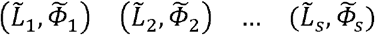, we similarly construct the kernelized meta score vector as 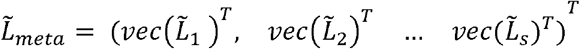 which under the null hypothesis follows a normal distribution with mean 0 and variance-covariance matrix 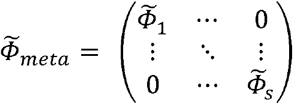. Then the test statistics can be constructed as 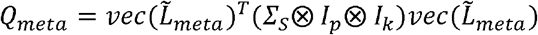 which follows a mixture of chi-square under the null hypothesis. The mixing parameters of the null distribution are the eigen values of 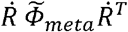 where 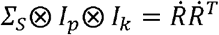.

## Section C: Resampling algorithm

In equation (4), *L_s_ ~ N*(0, *Φ_S_*) and in particular *L_meta_ ~ N*(0, *Φ_meta_*) under the null hypothesis of no association. Suppose we have B Meta-MultiSKAT tests with corresponding kernel matrices *Σ_S_, Σ_P_* and *Σ_G_* and the corresponding p-values T_P_ = (p_1_, p_2_, …, p_B_). Our test statistic is p_min_ = min(p_1_, p_2_, …, p_B_). Here we adopt a resampling based approach to estimate this correlation structure.

1. Generate null *L_s;null_ ~ N*(0, *Φ_s_*) for s = 1,2,…,S and form *L_meta;null_*
2. Calculate 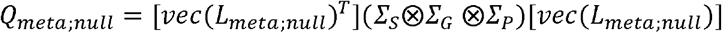 for each combination of *Σ_S_, Σ_P_* and *Σ_G_*
3. Calculate the corresponding asymptotic p-value
4. Repeat the previous steps for R (= 500 or 1000) times and calculate the null correlation between the p-values

With the estimated null correlation structure, we use a t-Copula to approximate the joint distribution of T_P_. The final p-value for p_min_ is then calculated from the distribution function of the assumed t-Copula. As the same way, the resampling approach can be used with the kernelized score 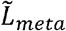.

## Section D: Illustration: Missing phenotype

To demonstrate the utility of Meta-MultiSKAT to handle missing phenotypes, we performed an analysis with all the 4 WBC phenotypes (lymphocyte, monocyte, basophil, eosinophill) in SardiNIA and only 3 (monocyte, basophil and eosinophil) in MGI. The models used for individual studies to extract the summary statistics remained the same.

We used Meta-MultiSKAT-Common-Rare tests in this analysis (Supplemental Table S2). All the variants, common and rare, were used in this analysis. The genes that were identified in the previous analysis were found to be significant or suggestive (p-value < 10^−5^) in this analysis as well, but with slightly differing p-values. As before, *PRG2, RP11-872D17*.8, IRF8 and CCL24 found to be significant using Meta-MultiSKAT methods.

## References

Astle, W. J., Elding, H., Jiang, T., Allen, D., Ruklisa, D., Mann, A. L., … Soranzo, N. (2016). The Allelic Landscape of Human Blood Cell Trait Variation and Links to Common Complex Disease. Cell, 167(5), 1415–1429.e19. https://doi.org/10.1016/j.cell.2016.10.042

Broadaway, K. A., Cutler, D. J., Duncan, R., Moore, J. L., Ware, E. B., Jhun, M. A., … Epstein, M. P. (2016). A Statistical Approach for Testing Cross-Phenotype Effects of Rare Variants. The American Journal of Human Genetics, 98(3), 525–540. https://doi.org/10.1016/j.ajhg.2016.01.017

Bulik-Sullivan, B. K., Loh, P.-R., Finucane, H. K., Ripke, S., Yang, J., Patterson, N., … Neale, B. M. (2015). LD Score regression distinguishes confounding from polygenicity in genome-wide association studies. Nature Genetics, 47(3), 291–295. https://doi.org/10.1038/ng.3211

Cotsapas, C., Voight, B. F., Rossin, E., Lage, K., Neale, B. M., Wallace, C., … Daly, M. J. (2011). Pervasive Sharing of Genetic Effects in Autoimmune Disease. PLoS Genetics, 7(8), e1002254. https://doi.org/10.1371/journal.pgen.1002254

Crosslin, D. R., McDavid, A., Weston, N., Zheng, X., Hart, E., de Andrade, M., … Jarvik, G. P. (2013). Genetic variation associated with circulating monocyte count in the eMERGE Network. Human Molecular Genetics, 22(10), 2119–2127. https://doi.org/10.1093/hmg/ddt010

Dahl, A., Iotchkova, V., Baud, A., Johansson, Å., Gyllensten, U., Soranzo, N., … Marchini, J. (2016). A multiple-phenotype imputation method for genetic studies. Nature Genetics, 48(4), 466–472. https://doi.org/10.1038/ng.3513

De Jager, P. L., Jia, X., Wang, J., de Bakker, P. I. W., Ottoboni, L., Aggarwal, N. T., … Oksenberg, J. R. (2009). Meta-analysis of genome scans and replication identify CD6, IRF8 and TNFRSF1A as new multiple sclerosis susceptibility loci. Nature Genetics, 41(7), 776–782. https://doi.org/10.1038/ng.401

Demarta, S., & McNeil, A. J. (2007). The t Copula and Related Copulas. International Statistical Review, 73(1), 111–129. https://doi.org/10.1111/j.1751-5823.2005.tb00254.x

Dutta, D., Scott, L., Boehnke, M., & Lee, S. (2019). Multi-SKAT: General framework to test for rare-variant association with multiple phenotypes. Genetic Epidemiology, 43(1), 4–23. https://doi.org/10.1002/gepi.22156

Engel, S. M., Molldrem, J. J., Lee, S., Harmon, Q. E., Wu, M. C., Armistead, P. M., … Maity, A. (2013). Kernel Machine SNP-Set Testing Under Multiple Candidate Kernels. Genetic Epidemiology, 37(3), 267–275. https://doi.org/10.1002/gepi.21715

Erlinger, T. P., Muntner, P., & Helzlsouer, K. J. (2004). WBC count and the risk of cancer mortality in a national sample of U.S. adults: results from the Second National Health and Nutrition Examination Survey mortality study. Cancer Epidemiology, Biomarkers & Prevention□: A Publication of the American Association for Cancer Research, Cosponsored by the American Society of Preventive Oncology, 13(6), 1052–1056. Retrieved from http://www.ncbi.nlm.nih.gov/pubmed/15184263

Ferreira, M. A. R., & Purcell, S. M. (2009). A multivariate test of association. Bioinformatics, 25(1), 132–133.

Fritsche, L. G., Gruber, S. B., Wu, Z., Schmidt, E. M., Zawistowski, M., Moser, S. E., … Mukherjee, B. (2018). Association of Polygenic Risk Scores for Multiple Cancers in a Phenome-wide Study: Results from The Michigan Genomics Initiative. The American Journal of Human Genetics, 102(6), 1048–1061. https://doi.org/10.1016/j.ajhg.2018.04.001

Hasegawa, T., Negishi, T., & Deguchi, M. (2002). WBC Count, Atherosclerosis and Coronary Risk Factors. Journal of Atherosclerosis and Thrombosis, 9(5), 219–223. https://doi.org/10.5551/jat.9.219

He, Z., Xu, B., Lee, S., & Ionita-Laza, I. (2017). Unified Sequence-Based Association Tests Allowing for Multiple Functional Annotations and Meta-analysis of Noncoding Variation in Metabochip Data. The American Journal of Human Genetics, 101(3), 340–352. https://doi.org/10.1016/j.ajhg.2017.07.011

Huang, J., Johnson, A. D., & O’Donnell, C. J. (2011). PRIMe: a method for characterization and evaluation of pleiotropic regions from multiple genome-wide association studies. Bioinformatics, 27(9), 1201–1206. https://doi.org/10.1093/bioinformatics/btr116

Ionita-Laza, I., Capanu, M., De Rubeis, S., McCallum, K., & Buxbaum, J. D. (2014). Identification of Rare Causal Variants in Sequence-Based Studies: Methods and Applications to VPS13B, a Gene Involved in Cohen Syndrome and Autism. PLoS Genetics, 10(12), e1004729. https://doi.org/10.1371/journal.pgen.1004729

Ionita-Laza, I., Lee, S., Makarov, V., Buxbaum, J. D., & Lin, X. (2013). Sequence Kernel Association Tests for the Combined Effect of Rare and Common Variants. The American Journal of Human Genetics, 92(6), 841–853. https://doi.org/10.1016/j.ajhg.2013.04.015

Kanai, M., Akiyama, M., Takahashi, A., Matoba, N., Momozawa, Y., Ikeda, M., … Kamatani, Y. (2018). Genetic analysis of quantitative traits in the Japanese population links cell types to complex human diseases. Nature Genetics, 50(3), 390–400. https://doi.org/10.1038/s41588-018-0047-6

Keller, M. F., Reiner, A. P., Okada, Y., van Rooij, F. J. A., Johnson, A. D., Chen, M.-H., … Nalls, M. A. (2014). Trans-ethnic meta-analysis of white blood cell phenotypes. Human Molecular Genetics, 23(25), 6944–6960. https://doi.org/10.1093/hmg/ddu401

Kim, D. K., Cho, M. H., Hersh, C. P., Lomas, D. A., Miller, B. E., Kong, X., … Silverman, E. K. (2012). Genome-Wide Association Analysis of Blood Biomarkers in Chronic Obstructive Pulmonary Disease. American Journal of Respiratory and Critical Care Medicine, 186(12), 1238–1247. https://doi.org/10.1164/rccm.201206-1013OC

Kim, J. H., Lim, S., Park, K. S., Jang, H. C., & Choi, S. H. (2017). Total and differential WBC counts are related with coronary artery atherosclerosis and increase the risk for cardiovascular disease in Koreans. PLoS ONE. https://doi.org/10.1371/journal.pone.0180332

Kircher, M., Witten, D. M., Jain, P., O’Roak, B. J., Cooper, G. M., & Shendure, J. (2014). A general framework for estimating the relative pathogenicity of human genetic variants. Nature Genetics, 46(3), 310–315. https://doi.org/10.1038/ng.2892

Kurotaki, D., Nakabayashi, J., Nishiyama, A., Sasaki, H., Kawase, W., Kaneko, N., … Tamura, T. (2018). Transcription Factor IRF8 Governs Enhancer Landscape Dynamics in Mononuclear Phagocyte Progenitors. Cell Reports, 22(10), 2628–2641. https://doi.org/10.1016/j.celrep.2018.02.048

Lee, S., Teslovich, T. M., Boehnke, M., & Lin, X. (2013). General Framework for Meta-analysis of Rare Variants in Sequencing Association Studies. The American Journal of Human Genetics, 93(1), 42–53. https://doi.org/10.1016/j.ajhg.2013.05.010

Lee, S., Won, S., Kim, Y. J., Kim, Y., Kim, B.-J., & Park, T. (2017). Rare variant association test with multiple phenotypes. Genetic Epidemiology, 41(3), 198–209. https://doi.org/10.1002/gepi.22021

Li, B., & Leal, S. M. (2008). Methods for Detecting Associations with Rare Variants for Common Diseases: Application to Analysis of Sequence Data. The American Journal of Human Genetics, 83(3), 311–321. https://doi.org/10.1016/j.ajhg.2008.06.024

Madsen, B. E., & Browning, S. R. (2009). A Groupwise Association Test for Rare Mutations Using a Weighted Sum Statistic. PLoS Genetics, 5(2), e1000384. https://doi.org/10.1371/journal.pgen.1000384

Majumdar, A., Haldar, T., Bhattacharya, S., & Witte, J. S. (2018). An efficient Bayesian meta-analysis approach for studying cross-phenotype genetic associations. PLOS Genetics, 14(2), e1007139. https://doi.org/10.1371/journal.pgen.1007139

Manichaikul, A., Mychaleckyj, J. C., Rich, S. S., Daly, K., Sale, M., & Chen, W. M. (2010). Robust relationship inference in genome-wide association studies. Bioinformatics, 26(22), 2867–2873. https://doi.org/10.1093/bioinformatics/btq559

Mc, W., S, L., T, C., Y, L., M, B., & X, L. (2011). Rare-variant association testing for sequencing data with the sequence kernel association test. American Journal of Human Genetics, 89, 82–93.

McLaren, W., Gil, L., Hunt, S. E., Riat, H. S., Ritchie, G. R. S., Thormann, A., … Cunningham, F. (2016). The Ensembl Variant Effect Predictor. Genome Biology, 17(1), 122. https://doi.org/10.1186/s13059-016-0974-4

Mu oz, A., Peng, Y., Chmiel, J. S., Margolick, J., Oishi, J., Samet, J. M., … Kingsley, L. (2012). Longitudinal Relation between Smoking and White Blood Cells. American Journal of Epidemiology, 144(8), 734–741. https://doi.org/10.1093/oxfordjournals.aje.a008997

Panagiotou, O. A., Willer, C. J., Hirschhorn, J. N., & Ioannidis, J. P. A. (2013). The Power of Meta-Analysis in Genome-Wide Association Studies. Annual Review of Genomics and Human Genetics, 14(1), 441–465. https://doi.org/10.1146/annurev-genom-091212-153520

Purcell, S. M., Smoller, J. W., Cotsapas, C., Solovieff, N., & Lee, P. H. (2013). Pleiotropy in complex traits: challenges and strategies. Nature Reviews Genetics, 14(7), 483–495. https://doi.org/10.1038/nrg3461

Ray, D., & Boehnke, M. (2018). Methods for meta-analysis of multiple traits using GWAS summary statistics. Genetic Epidemiology, 42(2), 134–145. https://doi.org/10.1002/gepi.22105

Ray, D., Pankow, J. S., & Basu, S. (2016). USAT: A Unified Score-Based Association Test for Multiple Phenotype-Genotype Analysis. Genetic Epidemiology, 40(1), 20–34. https://doi.org/10.1002/gepi.21937

Ried, J. S., Döring, A., Oexle, K., Meisinger, C., Winkelmann, J., Klopp, N., … Gieger, C. (2012). PSEA: Phenotype Set Enrichment Analysis--a new method for analysis of multiple phenotypes. Genetic Epidemiology, 36(3), 244–252. https://doi.org/10.1002/gepi.21617

Salcedo, T., Li, Y., Kreider, B. L., Li, H., Patel, V. P., Leung, K., … Thotakura, R. (2002). Molecular and Functional Characterization of Two Novel Human C-C Chemokines as Inhibitors of Two Distinct Classes of Myeloid Progenitors. The Journal of Experimental Medicine, 185(7), 1163–1172. https://doi.org/10.1084/jem.185.7.1163

Salomon, S., Guignant, C., Morel, P., Flahaut, G., Brault, C., Gourguechon, C., … Goëb, V. (2017). Th17 and CD24hiCD27+ regulatory B lymphocytes are biomarkers of response to biologics in rheumatoid arthritis. Arthritis Research & Therapy, 19(1), 33. https://doi.org/10.1186/s13075-017-1244-x

Schaffner,Stephen F, Foo,Catherine, Gabriel,Stacey, Reich,David, Daly,Mark J, & Altshuler,David. (2005). Calibrating a coalescent simulation of human genome sequence variation. Genome Research, 15(11), 1576–1583. https://doi.org/10.1101/gr.3709305

Sichien, D., Scott, C. L., Martens, L., Vanderkerken, M., Van Gassen, S., Plantinga, M., … Guilliams, M. (2016). IRF8 Transcription Factor Controls Survival and Function of Terminally Differentiated Conventional and Plasmacytoid Dendritic Cells, Respectively. Immunity. https://doi.org/10.1016/j.immuni.2016.08.013

Sidore, C., Busonero, F., Maschio, A., Porcu, E., Naitza, S., Zoledziewska, M., … Abecasis, G. R. (2015). Genome sequencing elucidates Sardinian genetic architecture and augments association analyses for lipid and blood inflammatory markers. Nature Genetics, 47(11), 1272–1281. https://doi.org/10.1038/ng.3368

Sivakumaran, S., Agakov, F., Theodoratou, E., Prendergast, J. G., Zgaga, L., Manolio, T., … Campbell, H. (2011). Abundant Pleiotropy in Human Complex Diseases and Traits. The American Journal of Human Genetics, 89(5), 607–618. https://doi.org/10.1016/j.ajhg.2011.10.004

Urrutia, E., Lee, S., Maity, A., Zhao, N., Shen, J., Li, Y., & Wu, M. C. (2015). Rare variant testing across methods and thresholds using the multi-kernel sequence kernel association test (MK-SKAT). Statistics and Its Interface, 8(4), 495–505. https://doi.org/10.4310/SII.2015.v8.n4.a8

Vacca, L., Scuteri, A., Mameli, C., Dei, M., Loi, P., Abecasis, G. R., … Lakatta, E. (2006). Heritability of Cardiovascular and Personality Traits in 6,148 Sardinians. PLoS Genetics, 2(8), e132. https://doi.org/10.1371/journal.pgen.0020132

Wang, K., Li, M., & Hakonarson, H. (2010). ANNOVAR: functional annotation of genetic variants from high-throughput sequencing data. Nucleic Acids Research, 38(16), el64-el64. https://doi.org/10.1093/nar/gkq603

White, J. R., Imburgia, C., Dul, E., Appelbaum, E., O’Donnell, K., O’Shannessy, D. J., … Sarau, H. M. (1997). Cloning and functional characterization of a novel human CC chemokine that binds to the CCR3 receptor and activates human eosinophils. Journal of Leukocyte Biology, 62(5), 667–675. https://doi.org/10.1002/jlb.62.5.667

Wu, B., & Pankow, J. S. (2016). Sequence Kernel Association Test of Multiple Continuous Phenotypes. Genetic Epidemiology, 40(2), 91–100. https://doi.org/10.1002/gepi.21945

Wu, M. C., Lee, S., Cai, T., Li, Y., Boehnke, M., & Lin, X. (2011). Rare-Variant Association Testing for Sequencing Data with the Sequence Kernel Association Test. The American Journal of Human Genetics, 89(1), 82–93. https://doi.org/10.1016/j.ajhg.2011.05.029

Yáñez, A., Ng, M. Y., Hassanzadeh-Kiabi, N., & Goodridge, H. S. (2015). IRF8 acts in lineage-committed rather than oligopotent progenitors to control neutrophil vs monocyte production. Blood, 125(9), 1452–1459. https://doi.org/10.1182/blood-2014-09-600833

Zhan, X., Zhao, N., Plantinga, A., Thornton, T. A., Conneely, K. N., Epstein, M. P., & Wu, M. C. (2017). Powerful Genetic Association Analysis for Common or Rare Variants with High-Dimensional Structured Traits. Genetics, 206(4), 1779–1790. https://doi.org/10.1534/genetics.116.199646

Zhou, X., & Stephens, M. (2014). Efficient Algorithms for Multivariate Linear Mixed Models in Genome-wide Association Studies. Nat Genet, 11(4), 407–409. https://doi.org/10.1038/ng.2310

Zhu, X., Feng, T., Tayo, B. O., Liang, J., Young, J. H., Franceschini, N., … Levy, D. (2015). Meta-analysis of correlated traits via summary statistics from GWASs with an application in hypertension. American Journal of Human Genetics, 96(1), 21–36. https://doi.org/10.1016/j.ajhg.2014.11.011

