## Supplemental for "Meta-MultiSKAT: Multiple phenotype meta-analysis for region-based association test"

**Supplementary Tables and Figures**

| Gene | Lymphocytes | | Monocyte | | Basophil | | Eosinophil | | Joint (Multi-SKAT) | |
| --- | --- | --- | --- | --- | --- | --- | --- | --- | --- | --- |
|  | MGI | SardiNIA | MGI | SardiNIA | MGI | SardiNIA | MGI | SardiNIA | MGI | SardiNIA |

***Rare variant test***

| *PRG2* | 0.12 | 0.51 | 0.67 | 0.54 | 0.12 | 0.09 | 0.02 | $\boldsymbol{3.7\times}$10^-7^ | 0.48 | 8.8$\boldsymbol{\times}$10^-7^ |
| --- | --- | --- | --- | --- | --- | --- | --- | --- | --- | --- |
| *RP11-872D17.8* | 0.23 | 0.77 | 0.58 | 0.91 | 0.16 | 0.11 | 0.01 | $\boldsymbol{5.9\times}$**10^-7^** | 0.80 | 3.1$\boldsymbol{\times}$**10^-6^** |
| *Combined effect of common and rare variants* | | | | | | | | | | |
| *PRG2* | 0.39 | 0.54 | 0.73 | 0.61 | 0.12 | 0.07 | 0.08 | $\boldsymbol{7.8\times}$**10^-7^** | 0.37 | **4.6**$\boldsymbol{\times}$**10^-7^** |
| *IRF8* | 0.83 | 0.01 | $\boldsymbol{1.0\times}$**10^-5^** | 3$.9\times$10^-5^ | 0.76 | 0.31 | 0.97 | 0.93 | 5.9$\times$10^-5^ | 3.5$\times$10^-4^ |
| *CCL24* | 0.67 | 0.92 | 0.08 | 0.39 | 0.03 | $4.3\times$10^-5^ | 9.1$\times$10^-5^ | 0.69 | 2.0$\times$10^-4^ | 1.2$\times$10^-3^ |
| *RP11-872D17.8* | 0.33 | 0.74 | 0.81 | 0.88 | 0.24 | 0.10 | 0.29 | $\boldsymbol{1.1\times}$**10^-6^** | 0.62 | **9.1**$\boldsymbol{\times}$**10^-6^** |
| *CADD-score weighting* | | | | | | | | | | |
| *PRG2* | 0.26 | 0.43 | 0.71 | 0.42 | 0.06 | 0.06 | 0.04 | $\boldsymbol{7.1\times}$**10^-7^** | 0.17 | **2.9**$\boldsymbol{\times}$**10^-8^** |
| *IRF8* | 0.61 | 0.24 | $1.7\times$10^-5^ | 1.9$\times$10^-4^ | 0.63 | 0.29 | 0.88 | 0.81 | 1.1$\times$10^-5^ | 3.8$\times$10^-3^ |
| *CCL24* | 0.48 | 0.76 | 0.12 | 0.16 | 0.07 | $1.0\times$10^-4^ | 8.9$\times$10^-5^ | 0.53 | 1.3$\times$10^-4^ | 1.2$\times$10^-3^ |
| *RP11-872D17.8* | 0.41 | 0.59 | 0.63 | 0.81 | 0.11 | 0.08 | 0.10 | $\boldsymbol{8.9\times}$**10^-6^** | 0.27 | **4.3**$\boldsymbol{\times}$**10^-6^** |

**Table S1**: Single phenotype and Multi-SKAT p-value for each of the 4 WBC subtypes in each of MGI and SardiNIA studies. For single-phenotype gene-based tests for rare-variants and CADD-score weighting, SKAT-O was used.

| **Gene** | **Meta-Het** | **Meta-Hom** | **Meta-Com** |
| --- | --- | --- | --- |
| *PRG2* | **2.4×10^−7^** | 7.6×10^−5^ | **4.1×10^−7^** |
| *IRF8* | 6.5×10^−4^ | **1.7×10^−6^** | **2.8×10^−6^** |
| *CCL24* | **4.8×10^−6^** | 8.2×10^−3^ | **8.9×10^−6^** |
| *RP11-872D17.8* | **8.4×10^−7^** | 4.1× 10^−4^ | **1.2×10^−6^** |

**Table S2**: Genes/regions identified by either of the Meta-MultiSKAT methods (Meta-Hom, Meta-Het or Meta-Com) in the example with missing phenotypes. The p-values < 10^−5^ were marked in bold.

| **α** | **Meta-Hom** | **Meta-Het** | **Meta-Com** |
| --- | --- | --- | --- |
| 1× 10^−5^ | 1.3× 10^−5^ | 1.3× 10^−5^ | 1.4× 10^−5^ |
| 1× 10^−4^ | 1.1× 10^−4^ | 1.1× 10^−4^ | 1.3× 10^−4^ |

**Table S3**: Estimated Type-1 error rates for Meta-MultiSKAT-Common-Rare tests.

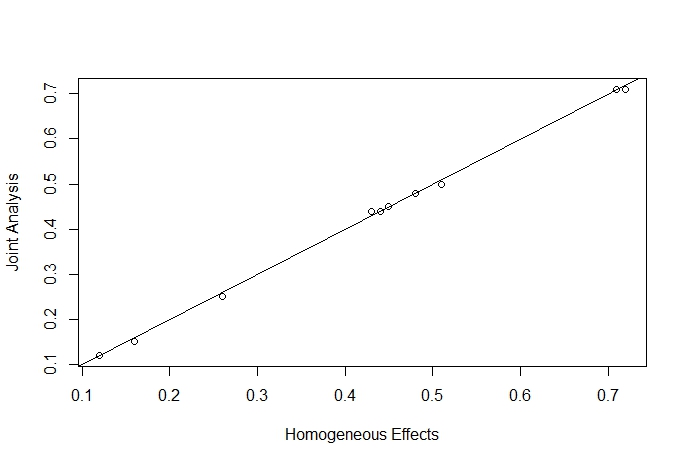

**Figure S1**: Power comparison for Joint analysis and analysis with $\Sigma_{S}= \Sigma_{S;Hom}$.

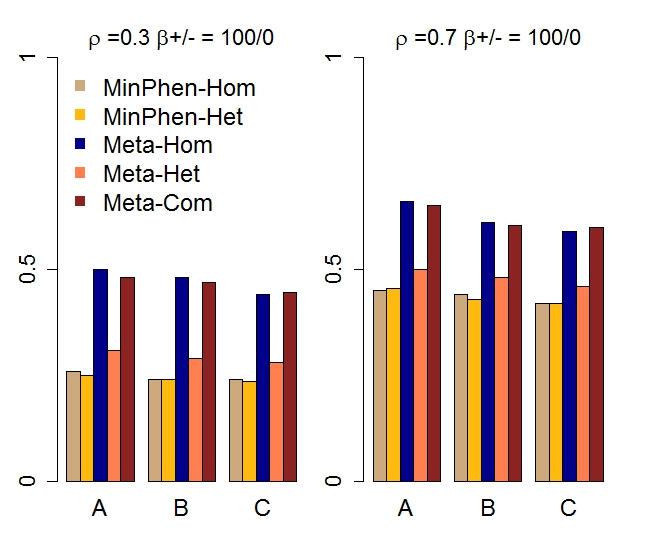

**Figure S2**: Power for Meta-MultiSKAT-Common-Rare tests compared with the existing methods when the set of causal variants is the same across different studies and has the same direction of effect. Empirical power for Meta-Hom, Meta-Het and Meta-Com plotted for 3 different scenarios compared against MinPhen-Hom and MinPhen-Het (See Simulations for details). Left panel shows the results for low correlation (ρ = 0.3) among the phenotypes and right panel shows the results for high correlation (ρ = 0.7).

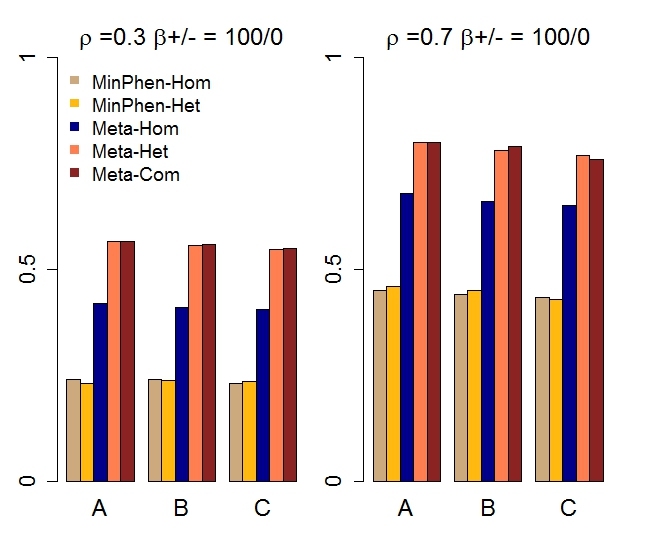

**Figure S3**: Power for Meta-MultiSKAT-Common-Rare tests compared with the existing methods when the set of causal variants is randomly chosen for each study and has the same direction of effect. Empirical power for Meta-Hom, Meta-Het and Meta-Com plotted for 3 different scenarios compared against MinPhen-Hom and MinPhen-Het (See Simulations for details). Left panel shows the results for low correlation (ρ = 0.3) among the phenotypes and right panel shows the results for high correlation (ρ = 0.7).

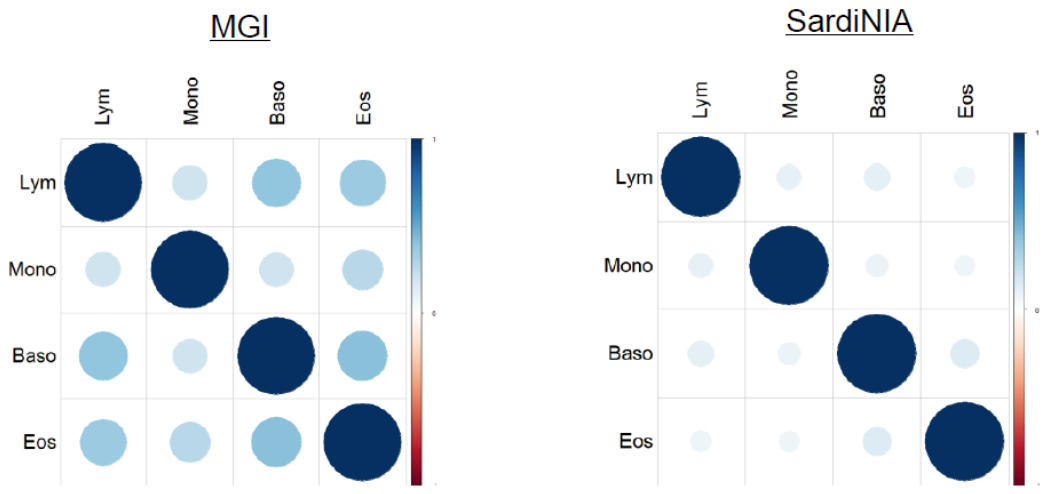

**Figure S4**: Correlation structure of the WBC phenotypes in MGI and SardiNIA respectively. Lym: Lymphocytes; Mono: Monocytes; Baso: Basophils; Eos:Eosinophils.
